# Refining interaction search through signed iterative Random Forests

**DOI:** 10.1101/467498

**Authors:** Karl Kumbier, Sumanta Basu, James B. Brown, Susan Celniker, Bin Yu

## Abstract

Advances in supervised learning have enabled accurate prediction in biological systems governed by complex interactions among biomolecules. However, state-of-the-art predictive algorithms are typically “black-boxes,” learning statistical interactions that are difficult to translate into testable hypotheses. The iterative Random Forest (iRF) algorithm took a step towards bridging this gap by providing a computationally tractable procedure to identify the stable, high-order feature interactions that drive the predictive accuracy of Random Forests (RF). Here we refine the interactions identified by iRF to explicitly map responses as a function of interacting features. Our method, signed iRF (s-iRF), describes “subsets” of rules that frequently occur on RF decision paths. We refer to these “rule subsets” as signed interactions. Signed interactions share not only the same set of interacting features but also exhibit similar thresholding behavior, and thus describe a consistent functional relationship between interacting features and responses. We describe stable and predictive importance metrics (SPIMs) to rank signed interactions in terms of their stability, predictive accuracy, and strength of interaction. For each SPIM, we define *null importance metrics* that characterize its expected behavior under known structure. We evaluate our proposed approach in biologically inspired simulations and two case studies: predicting enhancer activity and spatial gene expression patterns. In the case of enhancer activity, s-iRF recovers one of the few experimentally validated high-order interactions and suggests novel enhancer elements where this interaction may be active. In the case of spatial gene expression patterns, s-iRF recovers all 11 reported links in the gap gene network. By refining the process of interaction recovery, our approach has the potential to guide mechanistic inquiry into systems whose scale and complexity is beyond human comprehension.

## 1 Introduction

The complexity of biological interactions presents a substantial challenge for humans interested in extracting them from data. On the other hand, state-of-the-art supervised learning algorithms are well suited to learning complex, high-order, non-linear relationships in massive datasets as evidenced by their ability to accurately predict biological phenomena. For example, genome-wide maps of activity for a large number of regulatory factors in both humans and model organisms have already driven insights into the complex architecture of functional regulation [12, 6, 21]. Analyses of these extensive datasets have revealed the importance of dynamic, tissue-specific, high-order interactions that regulate diverse programs of gene expression across a variety of developmental and environmental contexts [32, 29, 18, 16].

Interactions in supervised learners provide models of how experimental manipulations may affect responses and candidate hypotheses for biological interactions. For instance, a supervised learner trained to predict enhancer activity may identify a collection of transcription factors (TFs) that are enriched among active enhancers. This information could be used to predict how manipulating these TFs would impact enhancer activity and to test whether learned interactions represent biological mechanisms or statistical associations. From this perspective, translating the complex interactions learned by predictive algorithms into human-exaplainable forms offers an invaluable opportunity to leverage large datasets to guide inquiry.

Here we propose an approach to extract explainable, rule-based interactions from an ensemble of decision trees by building on the iterative Random Forest algorithm (iRF) [2]. The interactions we identify are based on collections of simple rules of the form

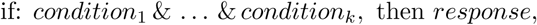

that define a clear functional relationship between interacting features and responses. As an example, conditions for continuous features **x** = (*x*_1_, …, *x_p_*) ∈ ℝ^*p*^ could take the form *x_j_* < *t_j_*, making rule-based interactions an attractive representation of the genomic processes that are activated by thresholded levels of regulatory factors [36]. In contrast to iRF, which searches for frequently co-occuring features on the decision paths of a Random Forest (RF) [4], the interactions we identify are “subsets” of rules that exhibit similar thresholding behavior across an RF. This representation provides additional information by describing both a precise relationship between interacting features and the subset of observations for which an interaction is active. Moreover, we find that the rule-based representation results in more accurate interaction recovery in our simulations.

To illustrate the benefit of our proposed approach, consider the following simple relationships between two regulatory factors and enhancer activity.

1. One subset of enhancers is activated by low concentrations of both factors while another subset is activated by high levels of both factors (Fig. 1 left).
2. Enhancers are activated by high concentrations of both factors (Fig. 1 middle)
3. Enhancers are actived by either high levels of either factor (Fig. 1 right).

**Figure 1:**
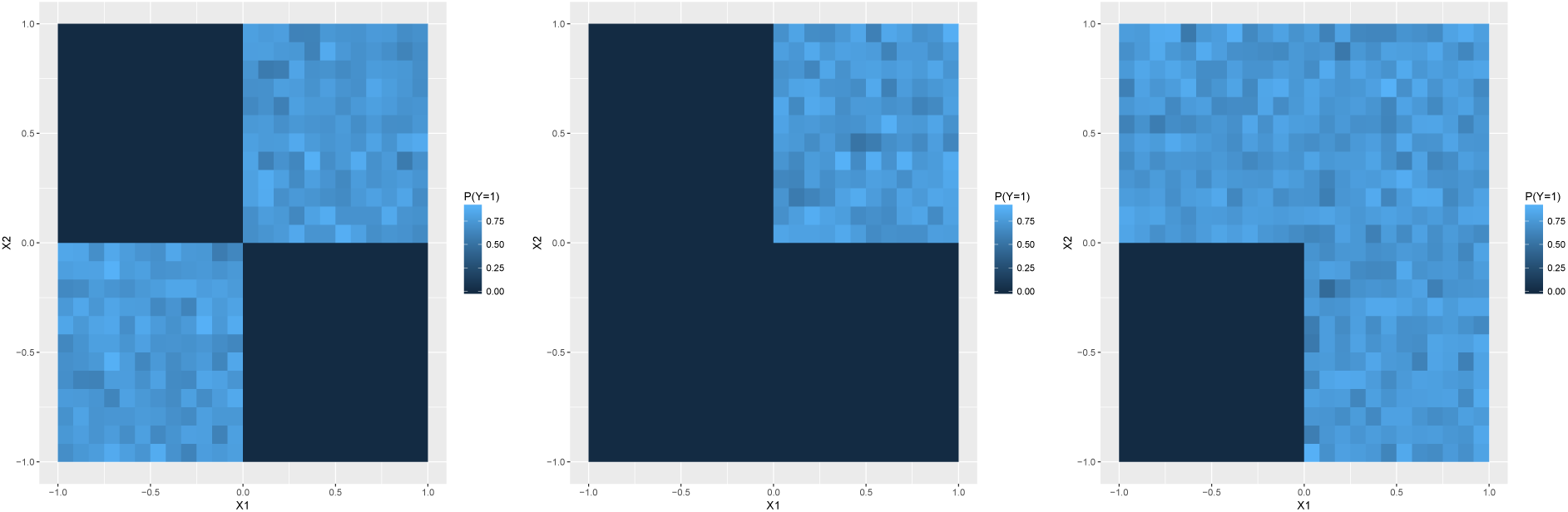
Proportion of active enhancers with responses generated from XOR (left), AND (middle), OR (right) rules. Each of these rules defines a unique functional relationship between regulatory factors and enhancer activity.

Enhancer activity in these examples always depends on the concentrations of both factors. However, each case defines a uniqe biological mechanism. Distinguishing between these settings is necessary to determine the experimental interventions that can reveal the underlying mechanism. iRF identifies that these regulatory factors influence enhancer activity. Our proposed approach distinguishes between these distinct settings and implicitly localizes these interactions to specific observations.

Current methods for extracting decision rules from data are largely based on decision trees, which partition observations into groups described by unique rules. Since rules learned by a single decision tree can be unstable with respect to small data perturbations [23], it is common to search for more reliable candidate interactions across ensembles of decision trees. One approach for identifying rules is to approximate an ensemble of decision trees through a single, stable tree [39, 17]. However, these methods rely on pseudo covariates, which can be difficult to generate in practice when the underlying data distribution is unknown. A second line of work extracts rules directly from an ensemble of decision trees: forest garrote [27], node harvest [28], rule fit [15], inTrees [7], and iRF [2]. With the exception of iRF, these methods rely on shallow trees to prevent overfitting and/or build rules through incremental search procedures that scale as *p^s^*, where *p* is the number of features and *s* is the order of an interaction. iRF scales independently of interaction order, enabling it to identify high-order interactions that arise in genomic applications. However, the interactions iRF recovers represent features whose collective activity is associated with responses, as opposed to a particular functional form.

Here we propose a refined iRF, signed iterative Random Forests (s-iRF), which describes the relationship between interacting features and responses. We use s-iRF to investigate regulatory interactions in the early stage *Drosophila* embryo. In particular, we study TF interactions that predict enhancer activity and that predict patterns of spatial gene expression. In the case of enhancer activity, we identify 15 of the 20 pairwise interactions reported in [2]. Three of the interactions we do not identify have either not been previously reported or are among TFs known to pattern different axes of the developing embryo. Additionally, we recover an interaction among gap genes *Giant (Gt), Krüppel (Kr*), and *Hunchback (Hb*), which is one of the few validated order-3 interactions in *Drosophila*. This interaction is known to regulate the *eve* stripe 2 enhancer, and our analysis suggests several other elements that it may regulate. In the case of spatial gene expression patterns, we recover all previously reported gap gene interactions and suggest new high-order interactions among anterior posterior (A-P) factors. Moreover, our predictive framing suggests that a large number of distinct interactions indicate positional information with high precision, pointing to a high level of redundancy in the A-P patterning system.

The rest of the paper is organized as follows. In section 2, we motivate the idea of signed interactions and introduce notations. In section 3, we present our three contributions:

- In section 3.1, we refine the interactions recovered by iRF through the notion of *signed interactions*, leading to signed iterative Random Forests (s-iRF). Signed interactions define a rule-based, functional relationship between interacting features and responses, and are thus more amenable to experimental follow-up. Moreover, the functional form of decision rules implicitly localizes signed interactions, describing the observations for which it is active.
- In section 3.2, we propose stable and predictive importance metrics (SPIMs) for signed interactions. These metrics expand upon the stability scores used by iRF, provding a more comprehensive assessment of the predictive accuracy, stability, and strength of an interaction.
- In section 3.3, we generalize the stability analysis framework of iRF to evaluate the uncertainty of our proposed metrics. In particular, we use *null importance metrics* that describe the expected behavior of our SPIMs when the data respect known structure to filter out observed results that can be reproduced by simple phenomena.

In section 4, we compare interactions recovered by iRF and s-iRF in simulations based on real and synthetic data. In section 5, we study signed interactions associated with enhancer activity and spatial gene expresion patterns in the early *Drosophila* embryo. We conclude by discussing areas for further work in interpreting complex supervised learning algorithms.

## 2 Background and notations

We motivate our problem with a simple example. Consider a collection of binary genomic responses *y_i_* ∈ {0, 1}, *i* = 1, …, *n*. For example, these responses may represent whether segment *i* of DNA is an enhancer in a particular context, with 0 indicating inactive and 1 active. For each response *i*, we measure a set of *p* features **x**_*i*_ = (*x*_*i*1_, …, *x_ip_*). These features typically represent the state, count, or concentration of biomolecules associated with segment *i*. For the sake of simplicity, suppose that we know whether each biomelecule is depleted, enriched, or within its normal range. This simplified framing is similar to [8], who binarize omics profiles based on divergence from baseline. In our simple setting, we may write **x**_*i*_ ∈ {−1, 0, 1}^*p*^. Of course, determining whether a biomolecule is enriched or depleted is challenging in practice. We will return to this issue shortly.

Given this collection of data we would like to identify configurations of the features, or interactions, for which there are a high proportion of active responses. More formally, we seek configurations *S* such that

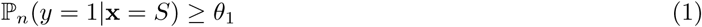

where ℙ_*n*_ denotes an empirical distribution and 0 ≤ *θ*_1_ ≤ 1. In the context of our example, equation (1) describes the enrichment of enhancer activity given the presence/absence of specific biomelecules. The enriched biomolecules (i.e. *S_j_* = 1) represent potential activators while the depleted features (i.e. *S_j_* = −1) represent potential repressors. Collectively, the features *j* such that *S_j_* ≠ 0 represent a potential order *s* = ∣*S*∣ interaction and a candidate biological mechanism.

One way to satisfy equation (1) is to identify highly specific interactions *S* that describe a very small number of observations. That is, interactions where ∣{*i*: **x**_*i*_ = *S*}∣ is small and ℙ_*n*_(*y* = 1∣**x** = *S*) is large. If ∣{*i*: **x**_*i*_ = *S*}∣ is particularly small, *S* may be a poor candidate for biological mechanisms that generalize beyond highly specific contexts. To ensure that the interactions represent a suitable proportion of active observations, we also seek interactions *S* such that

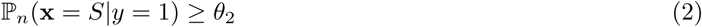

where 0 ≤ *θ*_2_ ≤ 1. From the perspective of our example, equation (2) describes the prevalence of an interaction *S* among all active enhancers. Of course, identifying local interactions that describe small subpopulations is of interest in some settings. The appropriate trade off between equations (1) and (2) is thus domain dependent.

In the next several sections we describe s-iRF, an approach for identifying and evaluating interactions *S* that describe which features are enriched, depleted, or within their normal range among a particular response class. We do so by extracting interactions from the decision paths of a Random Forest (RF) [4] through the iterative Random Forests (iRF) [2] framework. This allows us to identify high-order, rule-based interactions without forward-wise search procedures that scale exponentially in the size of interactions. The next section outlines the iRF algorithm. For a more detailed description, we refer readers to the original paper [2]

### 2.1 iterative Random Forests (iRF)

The iterative Random Forest algorithm provides a computationally efficient procedure to search for collections of interacting features that frequently co-occur along RF decision paths. This allows us to identify rules across an RF that share a common set of active features and may be good candidates for grouping into feature-restricted rules. We briefly review iRF here. For a full description of the algorithm, we refer readers to the original paper [2].

iRF trains a series of *K* iteratively re-weighted RFs, 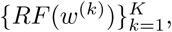 where features are sampled at each node with probability proportional to *w*^(*k*)^ ∈ 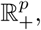 the Gini importance obtained from the previous iteration. Leaf node decision paths and predictions from the final iteration are encoded as pairs (𝓘_*i_t_*_, *Z_i_t__*), 𝓘_*i_t_*_ ⊆ {1, …, *p*}, *Z_i_t__* ∈ {0, 1}. 𝓘_*i_t_*_ denotes features selected along the decision path of the leaf node containing observation *i* = 1 …, *n* in tree *t* = 1, …, *T* and *Z_i_t__* the corresponding prediction. The pairs (𝓘_*i_t_*_, *Z_i_t__*) are used as input to RIT [33], which searches for frequently co-occuring features among a particular class of observations. The process of running RIT on (𝓘_*i_t_*_, *Z_i_t__*) produces a collection of feature interactions that frequently appear on the decision paths of RF(*w*^(*K*)^) and is described as generalized RIT (gRIT). Finally, iRF assess the stability of recovered interactions by repeating the gRIT step across an outer layer of bootstrap samples of the original data and evaluating the proportion of times an interaction is recovered.

## 3 Signed iterative Random Forests: an enhanced interpretation of iRF

We refine iRF interactions by searching for “rule subsets,” as opposed to selected features, that frequently occur along RF decision paths. Toward this end, we introduce *signed interactions*. Signed interactions formalize the notion of a rule subset and can be recovered efficiently from an RF. Intuitively, a signed interaction defines a decision path up to threshold. This allows us to compare decision paths whose thresholds vary as a result of the randomness introduced during RF training. Alternatively, a signed interaction can be viewed as a coarse region of the feature space corresponding to a predictive rule with smooth decision boundaries.

### 3.1 Signed interactions

Suppose that instead of data **x** ∈ {−1, 0, 1}^*p*^ we have continuous valued measurements **x** ∈ ℝ^*p*^. This represents a more realistic setting in practice, where we may not know *a priori* which features are enriched, depleted, or within their normal range. We map continuous features into the desired form using the decision paths of an RF, which can be represented as rules of the form

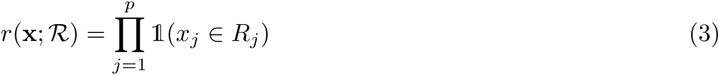

where 𝟙(⋅) denotes an indicator function, 𝓡 denotes a hyperrectangle associated with the decision path, and *R_j_* represent intervals of the form *x_j_* ≥ *t_j_* or *x_j_* < *t_j_*. For simplicity, we assume that each features is selected at most once along a decision path. We discuss how to handle other cases in 7.3.

To map continuous features **x** ∈ ℝ^*p*^ ↦ {−1, 0, 1}^*p*^, consider a decision tree node that splits on feature *j* with associated threshold *t_j_*. Observations **x** that arrive at this node will be sent to the left child when *x_j_* < *t_j_* and be sent to the right child otherwise. We use the signed feature-index *γ* ∈ {−*p*, …, *p*} to describe the selected splitting feature and inequality direction associated with a child node. That is, the left child is represented by the signed feature index *γ_ℓ_* = −*j* and the right child by *γ_r_* = *j* (Fig. 3). Under this definition the root node of a decision tree is not associated with a signed feature index. We define the *signed interaction* associated with leaf node *l* = 1, …, *L* as

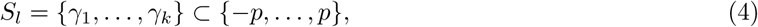

where *γ*_1_, …, *γ_k_* are the signed feature indices corresponding to all non-root nodes along the decision path (Fig. 2A). We note that there is a one-to-one correspondence between signed interactions defined in equation (4) and the interactions described in section 2. That is, we can write **x** ∈ {−1, 0, 1}^*p*^ as the signed interaction

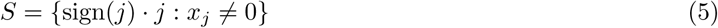

**Figure 2:**
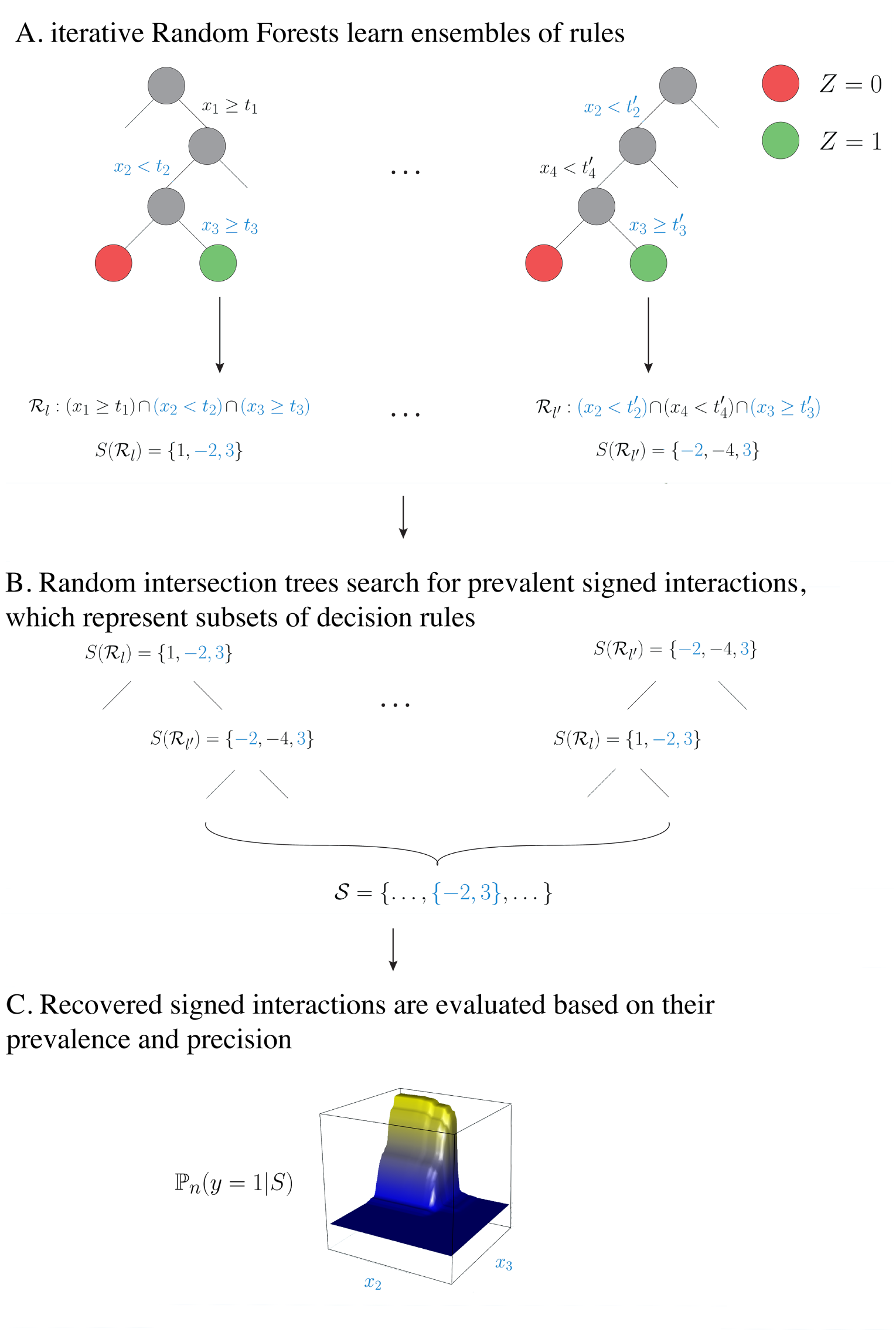
Mapping from RF decision rules to signed interactions. Leaf nodes that define hyperrectangles 𝓡 and 𝓡′ describe distinct regions of the feature space, but share the portion of their decision rule (highlighted in blue) up to threshold. (A) Each leaf node is encoded using a signed interaction (B) gRIT searches for prevalent signed interactions among RF decision paths (C) recovered signed interactions are evaluated based on importance metrics describing their stability and predictive accuracy.

**Figure 3:**
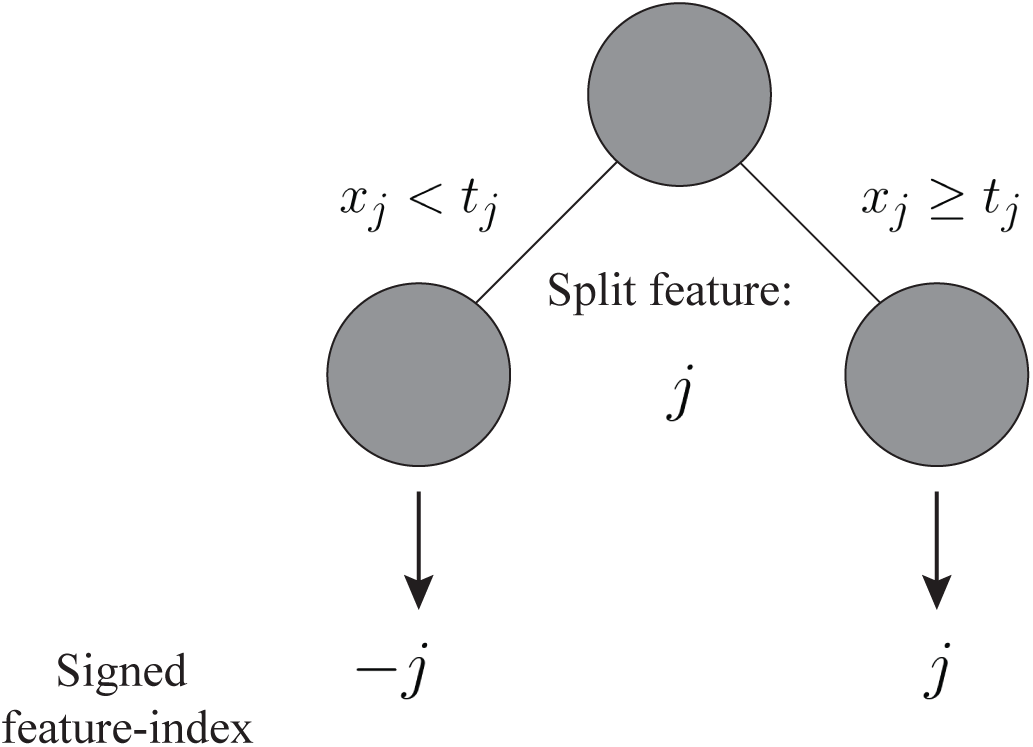
Decision tree directed feature mapping for node splits.

**Figure 4:**
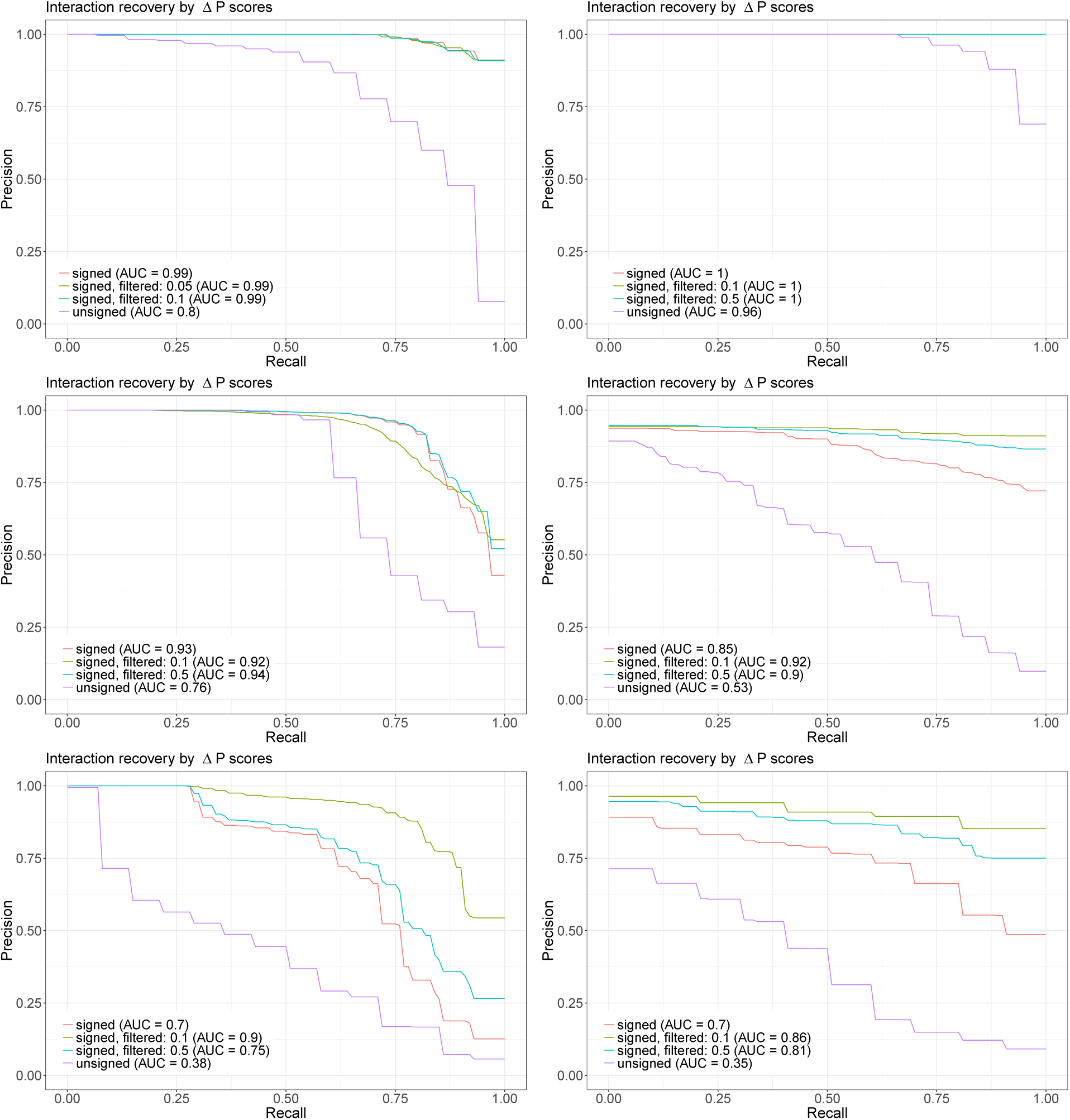
Simulation results for interaction recovery using *Drosophila* data (left) and standard Gaussian data (right) with responses generated from a single component AND rule (top), multi-component and rule (middle) and additive combination of and rules (bottom). PR curves for interaction recovery are given for unsigned interactions (purple), signed interactions (red), and signed interactions filtered based on PCS testing described in section 3.3 (blue, green).

Intuitively, signed interactions describe the rule associated with a decision path up to threshold. In other words, given a signed interaction *S* we can write

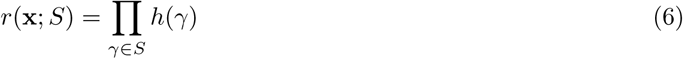

where

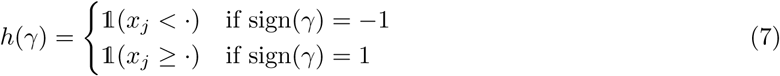

For instance, rules of the form 𝟙(*x*_2_ < ⋅)⋅𝟙(*x*_3_ ≥ ⋅) are represented by the signed interaction {−2, 3}. A signed interactions allows us to describe 2^*s*^ rules (up to threshold) that could be defined over *s* interacting features. In the context of genomic data, the different rules may suggest candidates for activators or repressors of a particular cellular process (e.g. Fig. 1). From a practical perspective, identifying how responses vary with different configurations of features can drastically reduce the number of experiments required to identify interesting biological mechanisms.

#### 3.1.1 Extracting signed interactions through iRF

Signed interactions that frequently appear on RF decision paths correspond to decision rules that an RF consistently selects under data and model perturbations. We search for prevalent signed interactions using gRIT, iRF’s interaction search procedure. Specifically, we generate signed feature-index sets for every leaf node in an RF and represent each observation with *T* pairs of signed feature-index sets and responses

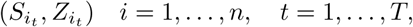

where *S_i_t__* represents the signed feature-index set for the leaf node containing observation *i* in tree *t* and *Z_i_t__* the corresponding prediction. These pairs serve as inputs to RIT [33]. Replacing feature-index sets with signed feature-index sets increases the number of regions gRIT searches over by a factor of 2^*s*^ while maintaining the same order of computational cost. This expressive representation allows us to determine more precise, and thus informative, regions of enriched response activity. In addition, we find in our simulations (section 4) that the signed representation improves the quality of recovered interactions. We call the process of searching for signed interactions using gRIT signed iRF (s-iRF).

### 3.2 Stable and predictive importance metrics

Here we define two importance metrics to evaluate signed interactions. Our first importance metric measures the stability of signed interactions based on their prevalence in a fitted RF. Our second importance metric evaluates the predictive accuracy of signed interactions based on response enrichment when the interaction is active. For each of these importance metrics, we define *null importance metrics* that serve as intuitive controls. Null importance metrics allow us to screen out signed interactions that can be explained by known, simple structure in the data. At a high level, our null metrics are inspired by [11], who consider the problem of generating controls for single neuron data.

#### 3.2.1 Prevalence to evaluate stability of signed interactions

Stability of results relative to “reasonable” data and model perturbations has been advocated as a minimum requirement to work towards reproducibility in science [38]. In the case of iRF, bootstrap sampling and random feature selection are natural perturbations to the data and model respectively. To evaluate the the stability of a signed interaction *S* relative to these perturbations, we consider its prevalence among class *C* ∈ {0, 1} leaf nodes

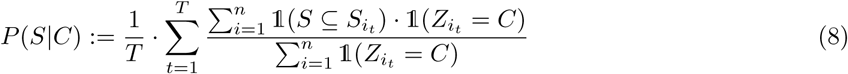

Prevalence indicates the proportion of class-*C* predicted observations where *S* is active, averaged across trees in an iRF. Equation (8) is identical to the notion of prevalence described in [33], and equation (2), except that it is defined relative to an ensemble of decision trees instead of a single dataset. As result, equation (8) incorporates the data and model perturbations used to train an iRF.

We note that equation (8) is closely related to the stability scores proposed by [2]. Specifically, stability scores represent the proportion of times an interaction is recovered by gRIT. The probability that an interaction is recovered by gRIT depends directly on its prevalence [33]. Thus prevalence can be viewed as a more refined measure of the stability scores. An added benefit of using prevalence to evaluate interactions is that unlike stability scores, it does not depend on RIT tuning parameters.

##### Null importance metric 1: class difference in prevalence

Our first null importance metric evaluates whether a signed interaction is more prevalent among one class of leaf nodes than another. In particular, we define the *class difference in prevalence* as

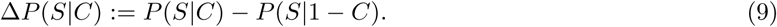

Equation (9) uses one class of leaf nodes as a control to determine whether a signed interaction is enriched among the other class. In our simulations, signed interactions can be recovered with high accuracy when ranked by Δ*P*(*S*∣1) (Fig. 6). In contrast, unsigned interactions identified by iRF include more false positives when ranked by Δ*P*(*S*∣ 1). Intuitively, the unsigned representation “washes out” important interactions when there is a high proportion of class-1 observations defined by one signed interaction (e.g. {2, 3}) and a high proportion of class-0 observations defined by a different signed interaction among the same features (e.g. {−2, 3}).

##### Null importance metric 2: independence of feature selection

Our second null importance metric evaluates the strength of a signed interaction. More precisely, it evaluates whether the features in a signed interaction are selected by an iRF in a dependent manner. For a given signed interaction *S*, consider the pairs 𝓟 = {(*S*′, *S*″): *S*′ ∪ *S*″ = *S*, *S*′ ∩ *S*″ = ∅, ∣*S*′∣ = 1}. We define the *independence of feature selection* as

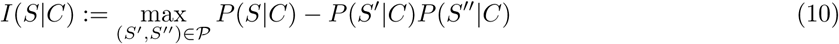

Equation (10) compares the prevalence of a signed interaction *S* with its expected prevalence if any single feature were selected independently of the others. In other words, it evaluates whether a set of features is collectively or individually associated with responses. From a modeling perspective, this is important because features often co-occur on decision paths after iterative re-weighting, regardless of whether they represent statistical interactions. From a biological perspective, this is particularly useful for identifying non-additive effects that underlie important natural processes. In our simulations, we find that *I*(*S*∣*C*) helps differentiate between additive and non-additive components of a generative model (Fig. 6).

#### 3.2.2 Precision to evaluate the predictive accuracy of signed interactions

Predictive accuracy quantifies how well a model emulates nature’s data generating process. Since signed interactions correspond to a subset of leaf nodes, we can associate them with a measure of predictive accuracy. More specifically, we evaluate a signed interaction *S* based on the distribution of responses among the subset of observations that appear in these leaf nodes. Formally, we define the *precision* of a signed interaction *S* for a class *C* as

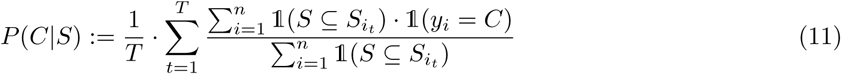

Of course, it might be desirable to evaluate an interaction based on other accuracy metrics. We describe an approach for extracting rule-based predictions associated with a signed interaction in 7.1. The predictions described in 7.1 can be evaluated using any metric of interest. Equation (11) represents a simple measure of accuracy that can be directly evaluated from a fitted iRF. To prevent inflated measures of accuracy due to overfitting, we report all measures of precision on held-out test data.

##### Null importance metric 3: increase in precision

Our final null metric evaluates whether a signed interaction is more precise than lower order subsets. We define the *increase in precision* as

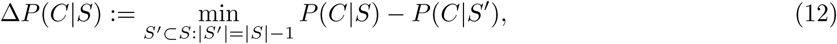

Intuitively, equation (12) evaluates whether all features in a signed interaction are necessary to achieve the observed level of precision. This allows us to identify the minimum set of features required to produce a level of predictive accuracy.

### 3.3 Generalized stability analysis to filter interactions

The importance measures defined in section 3.2 evaluate signed interactions recovered by a single s-iRF. To assess the uncertainty associated with these metrics, we evaluate them across RFs trained on an “outer layer” of bootstrap samples. This extends the stability analysis of iRF, which evaluates interactions recovered by gRIT across an outer layer of bootstrap samples [2]. That is, the importance metric in iRF is simply an indicator function specifying whether an interaction is recovered for each bootstrap sample *b* = 1, …, *B*.

In our generalized setting, the null metrics in section 3.2 describe attributes of an interaction relative to simple baselines. If any of Δ*P*(*S*∣*C*), *I*(*S*∣*C*), Δ*P*(*S*∣*C*) ≤ 0, the simple baseline provides a reasonable description of the observed data. Placing these null metrics within the iRF stability analysis framework allows us to quantify our certainty that

1. **Class enrichment**: *S* is more prevalent among class-1 leaf nodes than class-0 leaf nodes.
2. **Interaction strength**: Selection of features in S is driven by dependence in their joint distribution.
3. **Minimal representation**: All features in *S* are necessary for the observed level of prediction accuracy.

Algorithm 1 outlines our procedure to search for and filter interactions at a pre-defined level 0 ≤ *τ* ≤ 1. In our case studies, we apply algorithm 1 with *τ* = 0.5 to filter interactions recovered by s-iRF. This threshold improves the quality of recoered interactions in our simulation studies without being overly conservative. We are exploring the quality of screening under different generative models in our ongoing work.

## 4 Simulations

To evaluate the signed interactions recovered by iRF, we developed a suite of six simulation experiments. In particular, we generated responses from *n* = 1000 independent, standard Gaussian features **x** = (*x*_1_, …, *x*_50_) and *n* = 7808 *Drosophila* ChIP-chip measurements using three distinct generative models. For each simulation we drew responses *y* ~ *Bernoulli*(*π*), with *π* determined by Boolean rules intended to reflect the stereospecific interactions of biomolecules. We denote *π*^(*G*)^,*π*^(*D*)^ as the probability of response activity for the Gaussian and *Drosophila* data in each model respectively.

For responses generated by a signed interaction *S*^∗^ ⊆ {−*p*, …, *p*}, we define *j* ∈ *S*^∗^ as active signed feature indices. Any signed interaction *S* ⊆ *S*^∗^ containing only active signed feature indices is defined as an active interaction. A signed interaction *S* ⊈ *S*^∗^ containing any inactive signed feature indices is defined as inactive. We use these definitions as a “class label” for each signed interaction recovered by iRF. To evaluate the quality of interactions recovered by iRF, we calculate the area under the precision recall curve (AUC-PR) for recovered interactions, where each interaction is scored using Δ*P*(*S*∣1). Rankings based on the other metrics led to worse performance in all simulation settings.

### Algorithm 1

Interaction filtering at level *τ*

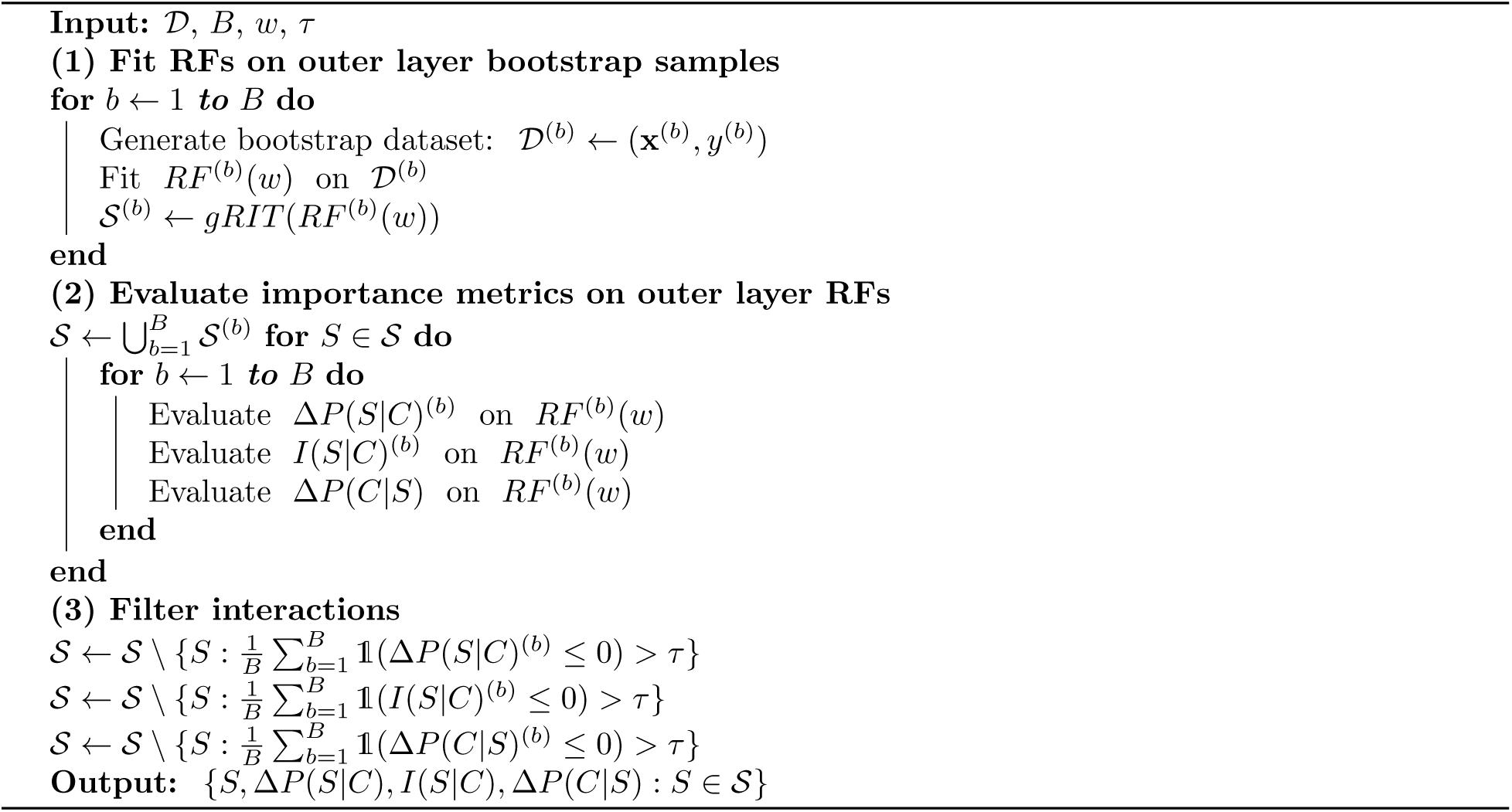

In each simulation experiment we ran iRF for 25 iterations over 100 bootstrap samples and report interactions recovered with *K* selected by maximizing prediction accuracy on out of bag samples. We grew 500 RITs with depth and number of children set to the default values of 5 and 2 respectively. For every setting, the precision recall curves for signed interactions show noticible improvement over the curves for unsigned interactions. When the generative model includes additive and non-additive components, the filtering procedure described in algorithm 1 further improves precision recall curves.

### 4.1 Single component AND rule

Our first model generates responses based on a single AND rule defined by high high concentrations of 4 features. We generated Bernoulli responses based on

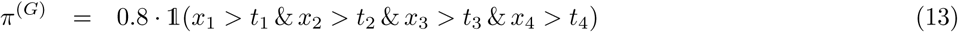

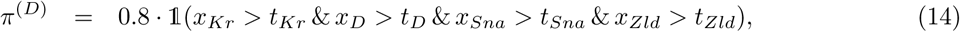

and set *t_j_* = *qt*(*x_j_*, 1−0.1^1/4^), which resulted in comparable class balance the true responses in the *Drosophila* data.

Fig. 6 (top) shows the precision-recall curves of interaction recovery based on Δ*P* scores for both the *Drosophila* (left) and Gaussian (right) data. In each plot, we report curves for unsigned interactions, signed interactions, and signed interactions filtered by the testing procedure outlined in Algorithm 1, averaged over 100 simulation replicates. For independent Gaussian features all three approaches achieve near perfect interaction recovery. That is, Δ*P* scores of recovered active interactions almost always exceed those of recovered inactive interactions. Highly correlated features in the *Drosophila* data have a minor impact on the quality of recovered signed interactions and a larger effect on unsigned interactions. Specifically, AUC-PR ranges from 0.8 (unsigned) to 0.99 (signed, signed and filtered). This is due to the fact that sparser signed interactions filter out large, inactive interactions that appear in the unsigned case where distinct signed interactions are condensed into an identical feature-index set.

### 4.2 Multi-component AND rule

Our second model generates responses based on two distinct AND rules defined over the same set of 4 features. We generated Bernoulli responses as

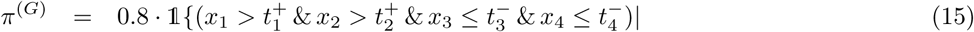

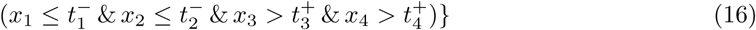

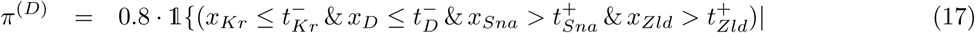

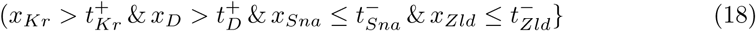

For the Gaussian data we set 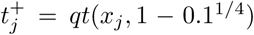 and 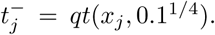 For the *Drosophila* data we set 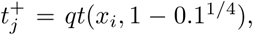 *j* ∈ {*Sna*, *Zld*}; 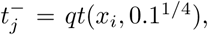 *j* ∈ {*D*, *Kr*}; 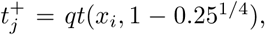 *j* ∈ {*D*, *Kr*}; 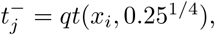 *j* ∈ {*Sna*, *Zld*}. The adjusted thresholds in the *Drosophila* data ensured a similar proportion of active responses for each and rule. In this setting, signed feature indices associated with the same AND rule were considered active interactions while those that were not were considered inactive. For instance, any combination of {1, 2, −3, −4} was considered an active interaction in the Gaussian setting while {1, 2, 3, 4} was considered inactive.

Fig. 6 (middle) shows the precision-recall curves for interaction recovery based on Δ*P* scores in both the *Drosophila* (left) and Gaussian (right) settings, averaged over 100 simulation replicates. Based on our definition of active interactions, there is a single active order-4 unsigned interaction among the 4 active features. There are 2^4^ signed interactions among these features, only two of which we consider active. Similarly, there are many inactive lower-order signed interactions that correspond to active unsigned interaction. Despite their more stringent definition, the quality of recovered signed interactions improves compared to unsigned interactions in both the Gaussian and *Drosophila* data. Once again, this is due to the fact that the signed representation implicitly filters out large, inactive interactions.

### 4.3 Additive AND rules

Our final model generates responses based on an additive combination of two AND rules defined over distinct sets of 3 features. We generated Bernoulli responses as

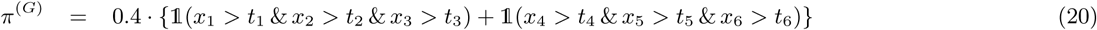

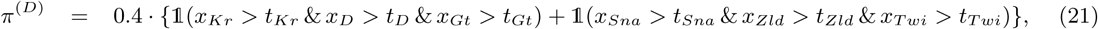

and set *t_j_* = *qt*(*x_j_*, 1−0.1^1/4^). In this setting, signed feature indices associated with the same AND rule were considered active interactions while that were not were considered inactive. For instance, any combination of {1, 2, 3} was considered an active interaction in the Gaussian setting while {1, 2, 4} was considered inactive.

Fig. 6 (bottom) shows the precision-recall curves for interaction recovery based on Δ*P* scores in both the *Drosophila* (left) and Gaussian (right) settings, averaged over 100 simulation replicates. Here iRF recovers many combinations of active features as unsigned interactions but does not distinguish between features that are participating in different AND rules. As a result, AUC-PR of unsigned interaction recovery drops considerably relative to previous settings for both the Gaussian and *Drosophila* data. On the other hand, the quality of recovered interactions improves substantially when we search among signed interactions, even when we do not explicitly filter out additive interactions using Algotrithm 1. The blue and green lines in Fig. 6 show curves for signed interaction recovery after removing interactions for which one of the restricted models achieves comparable predictive accuracy in more than 50% or 10% of bootstrap replicates respectively. By explicitly filtering interactions based on their form, we recover a set of signed interactions that both identify active features and distinguish between the functional relationship between those features and responses.

## 5 Searching for interactions in *Drosophila embryos*

Development and function in multicellular organisms depend on a precisely regulated programs of gene expression. Enhancers play a critical role in this process by coordinating the activity of TFs to increase transcription of target genes [5, 31]. In particular, distinct combinations of TFs bind enhancers to recruit regulatory co-factors, modify DNA structure, and/or interact directly with core promoter elements to regulate gene expression. Challenges associated with measuring the diverse set of genomic elements involved in these complex interactions make comprehensive, quantitative models of transcription inaccessible. Thus we considered simplified models to examine relationship between enhancers and TF activity measured through ChIP-chip and ChIP-seq assays as well as spatial gene expression images.

For both datasets, we represented candidate biological interactions as signed interactions, which are well suited to capture the combinatorial, thresholding behavior of well-studied biological processes [36, 9, 18]. We ran iRF for 25 iterations with 50 bootstrap samples and report interactions with *K* selected by minimizing prediction accuracy on out of bag samples. We grew 2500 RITs with number of children and depth set to default values of 2 and 5 respectively. Using many RITs allowed us to recover a large number of interactions that we subsequently filtered with algorithm 1.

### 5.1 ChIP assays and enhancer activity

Genome-wide, quantitative maps of DNA binding are available for 24 TFs known to play an important developmental role in early *Drosophila* embryos [26, 24]. These TFs bind to enhancer elements in distinct combinations to drive patterned gene expression, and subsequently embryogenesis. Based on experimental measurements of patterned gene expression, some 7809 genomic sequences have been labeled for enhancer activity in stage 5 blastoderm embryos [3, 13, 22]. Data collection and processing are described in [2].

To investigate potential regulatory interactions, we split the data into training and test sets of 3912 and 3897 genomic sequences respectively and trained iRF to predict enhancer activity (1: active element, 0: inactive element) of each sequence from TF binding levels. We filtered the recovered interactions using algorithm 1 with *τ* = 0.5. That is, all interactions we report are enriched among active enhancers (Δ*P*(*S*∣1) > 0), demonstrate evidence for interaction effects (*I*(*S*∣1) > 0), and are more precise than any lower order subset (Δ*P*(1∣*S*) > 0) in at least 50% of bootstrap replicates.

Table 1 reports the filtered signed interactions, ranked according to Δ*P*(*S*∣1). We recover 15 of the 20 pairwise interactions reported in [2]. These are indicated with ∗*s* in table 1. One of the interactions that we do not recover, among *Dichaete (D*) and *Twist (Twi*), has not been previously reported. Two of the other interactions we miss are among anterior posterior (A-P) and dorsal ventral (D-V) patterning factors, namely *Kr*, *Twi* and *Gt*, *Twi*. Although interactions among these elements have been previously reported [25], this evidence is weaker than for interactions among factors that belong to the same patterning system. These weaker interactions are precisely what we expect to filter when accounting for the strength of interaction using *I*(*S*∣1). It is interesting to note that all recovered interactions are defined by high binding levels TFs. In other words, all of our models reflect the notion that a TF must be bound to a genomic sequence somewhere in the embryo to represent a viable regulator of that element.

**Table 1:** Enhancer prediction signed interactions filtered at level *τ* = 0.5 and ranked according to Δ*P*(*S*∣1). Interactions indicated with ^∗^s represent the pairwise TF interactions reported in [2].

| Interaction | $\Delta P(S\vert 1)$ | $P(S\vert 1)$ | $P(1\vert S)$ |
| --- | --- | --- | --- |
| * Gt+; Zld+ | 0.218 | 0.224 | 0.602 |
| * Gt+; Kr+ | 0.204 | 0.21 | 0.543 |
| * Kr+; Zld+ | 0.172 | 0.193 | 0.506 |
| * Gt+; Hb+ | 0.154 | 0.157 | 0.576 |
| * Twi+; Zld+ | 0.149 | 0.183 | 0.437 |
| * Hb+; Zld+ | 0.128 | 0.134 | 0.544 |
| Gt+; Kr+; Zld+ | 0.117 | 0.119 | 0.64 |
| * Gt+; Med+ | 0.114 | 0.118 | 0.553 |
| * Hb+; Kr+ | 0.113 | 0.117 | 0.52 |
| Gt+; Hb+; Zld+ | 0.097 | 0.098 | 0.645 |
| * Kr+; Med+ | 0.096 | 0.105 | 0.492 |
| * Hb+; Twi+ | 0.095 | 0.1 | 0.487 |
| * Med+; Twi+ | 0.086 | 0.098 | 0.445 |
| Gt+; Hb+; Kr+ | 0.081 | 0.083 | 0.616 |
| * D+; Gt+ | 0.073 | 0.078 | 0.522 |
| Hb+; Kr+; Zld+ | 0.071 | 0.072 | 0.618 |
| D+; Kr+ | 0.063 | 0.08 | 0.44 |
| Gt+; Kr+; Med+ | 0.06 | 0.062 | 0.618 |
| Hb+; Med+ | 0.06 | 0.063 | 0.534 |
| D+; Zld+ | 0.058 | 0.083 | 0.433 |
| Hb+; Twi+; Zld+ | 0.057 | 0.058 | 0.596 |
| Dl+; Kr+ | 0.051 | 0.068 | 0.394 |
| Gt+; Hb+; Med+ | 0.043 | 0.044 | 0.622 |
| D+; Hb+ | 0.043 | 0.045 | 0.48 |
| D+; Med+ | 0.042 | 0.049 | 0.414 |
| Dl+; Zld+ | 0.04 | 0.077 | 0.367 |
| Dl+; Hb+ | 0.039 | 0.042 | 0.437 |
| Cad+; Gt+ | 0.02 | 0.02 | 0.561 |
| Cad+; Zld+ | 0.019 | 0.02 | 0.532 |
| * Bcd+; Zld+ | 0.019 | 0.02 | 0.495 |
| Cad+; Kr+ | 0.019 | 0.019 | 0.526 |
| Prd+; Twi+ | 0.018 | 0.02 | 0.394 |
| * Bcd+; Gt+ | 0.016 | 0.016 | 0.657 |
| * Bcd+; Twi+ | 0.016 | 0.016 | 0.459 |
| Med+; Prd+ | 0.012 | 0.013 | 0.438 |
| Cad+; Hb+ | 0.01 | 0.011 | 0.563 |
| Bcd+; Med+ | 0.01 | 0.011 | 0.523 |
| Cad+; Med+ | 0.01 | 0.01 | 0.49 |
| H+; Kr+ | 0.01 | 0.011 | 0.431 |
| H+; Med+ | 0.006 | 0.007 | 0.395 |
| Hb+; Sna+ | 0.004 | 0.004 | 0.525 |

**Table 2:** Experimentally validated enhancer elements with high predicted probability based on *Gt*, *Hb*, and *Kr* binding levels.

| ID | Neighboring genes | $g(\mathbf{x}; \{Gt+, Hb+, Kr+\})$ |
| --- | --- | --- |
| VT10589 | dia; cad | 1 |
| VT13456 | CR44446; CG44439 | 1 |
| VT14362 | CR43948; CG12134 | 1 |
| VT27678 | h; Pex7 | 1 |
| VT27680 | h; Pex7 | 0.999 |
| VT33934 | kni; cmpy | 0.999 |
| VT33935 | kni; cmpy | 0.999 |
| VT34803 | Ten-m; CG11449 | 0.999 |
| VT34804 | Ten-m; CG11449 | 0.998 |
| VT38550 | hb; Cks85A | 0.998 |
| VT55790 | gt; CG32797 | 0.992 |

In contrast to the interactions reported in [2], we identify several additional order-3 interactions (table 1). These concentrate around well-studied gap genes *Gt*, *Kr*, and *Hb* [20, 30, 10]. Of particular note, we identify an order-3 interaction among these factors, which suggests *Gt*, *Kr*, and *Hb* binding is enriched among active enhancers (Δ*P*(*S*∣1) = 0.081) and active enhancers are enriched ~ 6-fold when these three factors are bound (Δ*P*(*S*∣1) = 0.616). This is one of the few experimentally validated high-order TF interactions in the *Drosophila* embryo, having previously been studied in the case of *even*-*skipped* (*eve*) stripe 2 expression [34, 1]. Our results suggest that it is broadly important for establishing activity across other enhancers that have been less widely studied.

To identify candidate targets of an interaction among *Gt*, *Hb* and *Kr*, we generated predictions that depended exclusively on these three factors with the approach described in 7.1. Using these predictions, we considered the experimentally validated enhancer elements from [22] for which the predicted probability was > 0.99. Table 6 reports this subset of enhancers along with neighboring genes, which represent potential targets of the *Gt*, *Hb*, and *Kr* interaction. Of the 11 enhancers reported in table 6, 9 are annotated as driving A-P related expression patterns (either “A-P stripes” or “gap”). These represent well known gap gene targets in the early *Drosophila* embryo. Each of the remaining enhancers neighbor computational genes with unknown function, suggesting plausible novel targets of an interaction among *Gt*, *Hb*, and *Kr*.

In addition to well-studied gap gene interactions, the top four order-3 interactions are among TFs that are all known to interact in a pairwise manner. For instance, we identify the important interacting role of *Zelda (Zld*) in the early *Drosophila* embryo, particularly in the context of other gap genes. Fig. 5 depicts surface maps for an order-3 interaction among *Gt*, *Kr*, and *Zld*. When *Zld* binding is below the median RF threshold (left), the probability of a region acting as an active enhancer is as high as 0.23 if any two factors are bound at high levels but the third is bound at low levels. When all three factors are bound at high levels (right), the probability a region acts as an active enhancer is nearly 0.64, representing an approximately 3-fold increase. We note that this interaction was previously reported in [2] and is consistent with the belief that *Zld* acts as a potentiating factor that requires input from additional factors to regulate gene expression [14]. A similarly strong interaction among *Hb*, *Kr*, and *Zld* is shown in Fig. 6.

**Figure 5:**
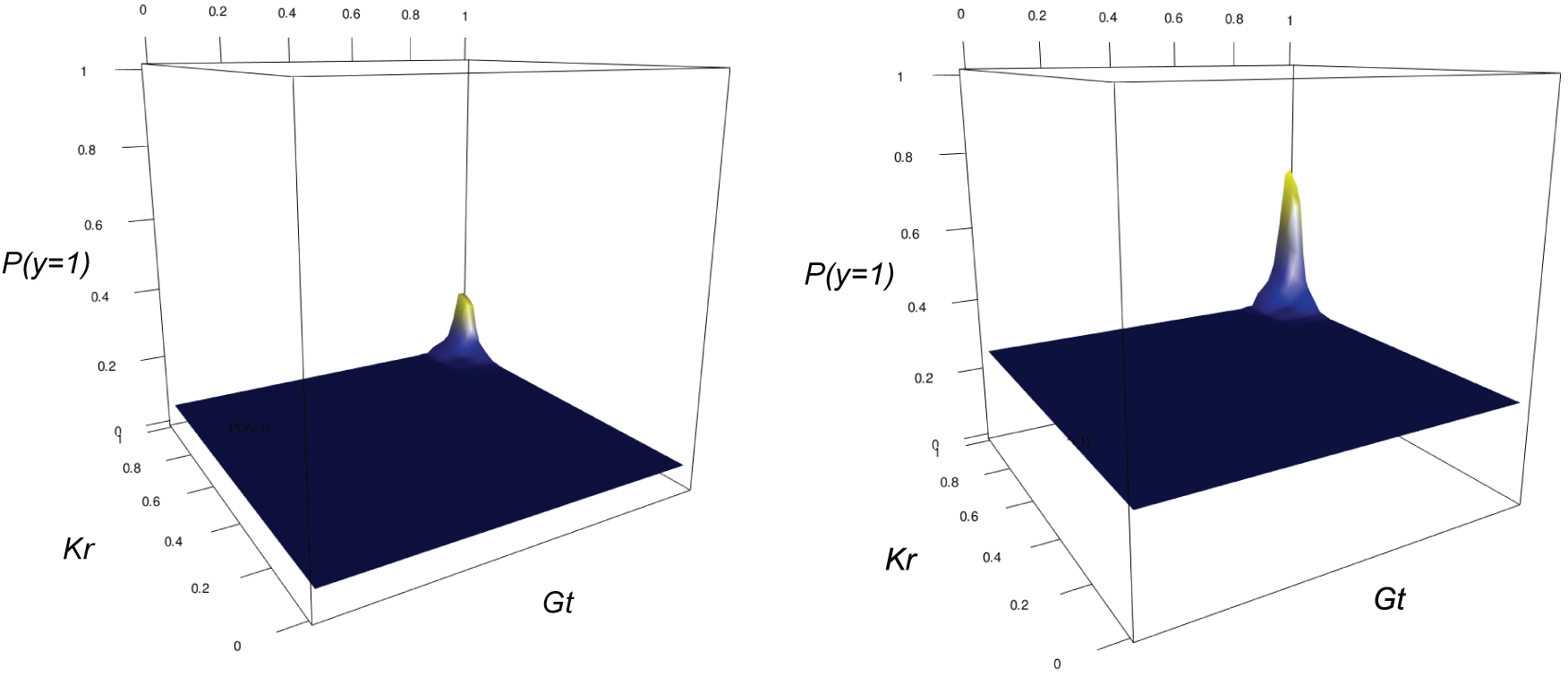
Surface maps depicting an order-3 interaction among Kr, Gt, and Zld. The probability that an observation acts as an active enhancer when all three factors are bound at high levels reaches as high as 0.64. When both Gt and Kr are bound at high levels but Zld is bound at low levels, the probability that an observation acts an active reaches as high as 0.23. This is indicative of a non-linear interaction where binding by all three factors represents nearly a three-fold increase in the probability of enhancer activity relative to binding by any two factors alone. Plots are drawn using held out test data to show the generalizability of this interaction.

**Figure 6:**
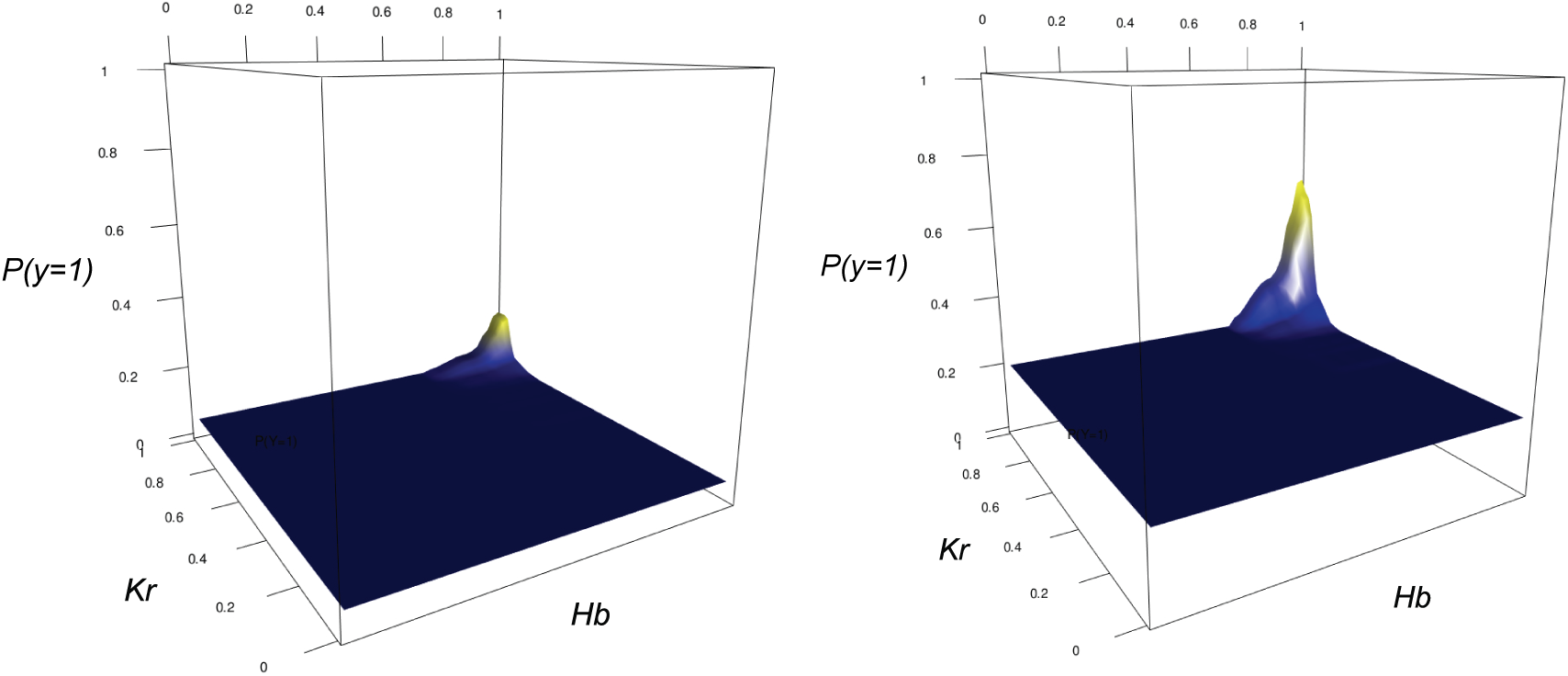
Surface maps depicting an order-3 interaction among Kr, Hb, and Zld. The probability that an observation acts as an active enhancer when all three factors are bound at high levels reaches as high as 0.62. When both Hb and Kr are bound at high levels but Zld is bound at low levels, the probability that an observation acts an active reaches as high as 0.21. This is indicative of a non-linear interaction where binding by all three factors represents nearly a three-fold increase in the probability of enhancer activity relative to binding by any two factors alone. Plots are drawn using held out test data to show the generalizability of this interaction.

**Figure 7:**
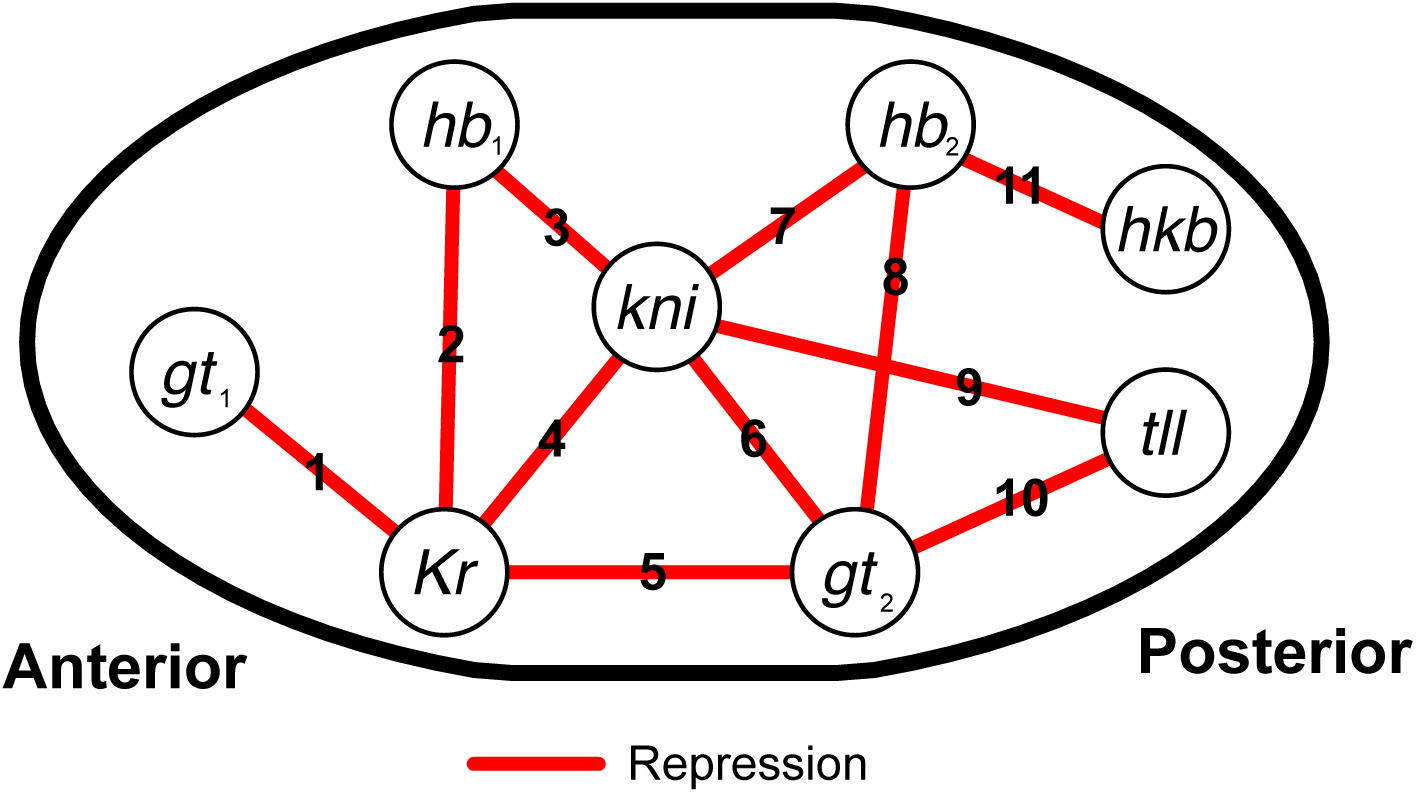
Spatial diagram of interactions in the gap gene network, as presented in [37] and originally described in [19]. Links are numbered 1 to 11 and all represent repressive interactions. Genes that are expressed in multiple, non-contiguous PPs are represented with subscripts (e.g. *gt*_1_, *gt*_2_).

### 5.2 Spatial gene expression and principal patterns

Studies of spatial gene expression have shown that complex expression patterns can be broken down into small regions regulated by enhancer elements and associated with organ primordia [37, 22, 34]. The *Berkeley Drosophila Genome Project* has generated over 137,211 gene expression images to study such spatial gene expression domains. These images represent localized gene expression in the *Drosophila* embryo measured through RNA *in situ* hybridization followed by confocal microscopy [35]. Previous work developed a quantitative representation of the images as 405 pixels to measure the concentration of gene expression in distinct regions of the embryo [37]. To condense this high dimensional data, the authors decompose these 405 regions into “principal patterns” (PPs) of gene expression. In particular, their spatial decomposition uses stability-driven nonnegative matrix factorization to represent a gene expression image as

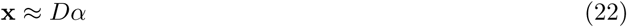

where 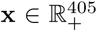 represents the processed image, 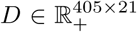 a dictionary of 21 learned PPs, and 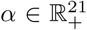 the PP coefficients for the given gene.

The intimate connection between enhancers and spatial gene expression make PPs a useful conceptual tool to relate spatial covariability of TFs to enhancer activity. We treated PPs as responses that define domains of gene expression and asked which TF interactions accurately reproduce these domains. Data acquisition and preprocessing are described in [37]. We split the data into randomly sampled training and test sets of 300 and 105 pixels respectively. Following [37], we defined a pixel as active for PP *i* if the corresponding dictionary atom *D*[, *i*] ≥ 0.1. With this definition, we generated responses for the 5 A-P segmentation stripe PPs (PP7-PP9, PP17, PP20), which we predicted using the spatial expression patterns of the TFs studied in section 5.1. This resulted in a total of 5 prediction problems. For each prediction problem, we trained an iRF to predict active pixels, and filtered interactions using the procedure describe in algorithm 1 at a level *τ* = 0.5.

Tables 3–7 report the interactions recovered from each PP prediction problem. These interactions largely surround gap and pair-ruled genes that are known play an important role in establishing the Δ*P* domain. Interestingly, many of the interactions we recover can predict a PP domain with high precision and are highly prevalent within the domain. For instance, almost all of the interactions we recover have precision *P*(1∣*S*) ≥ 0.8 and many have prevalence *P*(*S*∣1) > 0.2. In other words, a large number of distinct interactions can indicate positional information with high precision. This finding points to a high level of redundancy in the A-P patterning system.

**Table 3:** PP7 interactions, filtered at level *τ* = 0.5 and ranked according to Δ*P*(*S*∣1).

| Interaction | $\Delta P(S\vert 1)$ | $P(S\vert 1)$ | $P(1\vert S)$ |
| --- | --- | --- | --- |
| Kni-;Prd+ | 0.529 | 0.575 | 0.874 |
| Hb+;Prd+ | 0.446 | 0.47 | 0.902 |
| Cad-;Hb+ | 0.429 | 0.443 | 0.93 |
| Hb+;Kni-;Prd+ | 0.396 | 0.408 | 0.921 |
| Hb+;Run+ | 0.356 | 0.369 | 0.927 |
| Cad-;Run+ | 0.329 | 0.342 | 0.915 |
| Ftz+;Kni- | 0.313 | 0.334 | 0.896 |
| Prd+;Run+ | 0.311 | 0.328 | 0.914 |
| Cad-;Ftz+ | 0.285 | 0.29 | 0.932 |
| Cad-;Hb+;Run+ | 0.264 | 0.265 | 0.957 |
| Hb+;Prd+;Run+ | 0.245 | 0.249 | 0.942 |
| Cad-;Ftz+;Kni- | 0.233 | 0.235 | 0.939 |
| Hb+;Kni-;Prd+;Run+ | 0.206 | 0.208 | 0.953 |
| Kni-;Slp+ | 0.071 | 0.08 | 0.847 |
| Prd+;Slp- | 0.046 | 0.05 | 0.874 |
| Run+;Slp+ | 0.046 | 0.048 | 0.89 |
| Kni-;Prd+;Slp- | 0.043 | 0.046 | 0.886 |
| Kni-;Slp- | 0.042 | 0.053 | 0.842 |
| Hb+;Slp- | 0.041 | 0.044 | 0.896 |
| Ftz+;Slp+ | 0.04 | 0.041 | 0.905 |
| Hb+;Kni-;Slp- | 0.038 | 0.039 | 0.916 |
| Hb+;Prd+;Slp- | 0.038 | 0.039 | 0.92 |
| h+;Kni- | 0.034 | 0.037 | 0.849 |
| Kni-;Kr- | 0.019 | 0.022 | 0.835 |
| Cad-;Kr- | 0.015 | 0.016 | 0.865 |
| Kr-;Prd+ | 0.014 | 0.017 | 0.87 |
| Kr-;Run+ | 0.012 | 0.013 | 0.891 |
| Gt+;Kni- | 0.011 | 0.012 | 0.871 |
| D-;Kni- | 0.01 | 0.011 | 0.842 |
| D-;Prd+ | 0.009 | 0.01 | 0.856 |
| Hb+;Kr+ | 0.007 | 0.007 | 0.924 |

**Table 4:** PP8 interactions, filtered at level *τ* = 0:5 and ranked according to Δ*P*(*S*∣1).

| Interaction | $\Delta P(S\vert 1)$ | $P(S\vert 1)$ | $P(1\vert S)$ |
| --- | --- | --- | --- |
| D+;Kr+ | 0.143 | 0.145 | 0.99 |
| Gt-;Prd+ | 0.117 | 0.125 | 0.847 |
| Ftz+;Gt- | 0.111 | 0.118 | 0.889 |
| D+;Gt- | 0.077 | 0.085 | 0.842 |
| Cad-;Gt- | 0.075 | 0.082 | 0.864 |
| Gt-;h+ | 0.073 | 0.077 | 0.86 |
| Gt-;Run+ | 0.073 | 0.082 | 0.837 |
| Cad+;Gt- | 0.054 | 0.065 | 0.842 |
| Gt-;Hb- | 0.049 | 0.052 | 0.864 |
| Cad+;h+ | 0.027 | 0.035 | 0.825 |
| Cad-;Prd- | 0.016 | 0.019 | 0.798 |
| Shn-;Slp+ | 0.005 | 0.007 | 0.787 |
| D-;Kni+ | 0.002 | 0.004 | 0.78 |

**Table 5:** PP9 interactions, filtered at level *τ* = 0.5 and ranked according to Δ*P*(*S*∣1).

| Interaction | $\Delta P(S\vert 1)$ | $P(S\vert 1)$ | $P(1\vert S)$ |
| --- | --- | --- | --- |
| Cad+;Kni+ | 0.473 | 0.483 | 0.901 |
| Ftz+;Kni+ | 0.222 | 0.226 | 0.914 |
| D+;Kni+ | 0.112 | 0.114 | 0.899 |
| Cad+;h+ | 0.105 | 0.109 | 0.893 |
| Cad+;D+ | 0.103 | 0.109 | 0.852 |
| Cad+;D+;Kni+ | 0.081 | 0.082 | 0.909 |
| Cad+;Prd+ | 0.078 | 0.082 | 0.884 |
| D+;Hb- | 0.073 | 0.077 | 0.859 |
| Cad+;Tll- | 0.065 | 0.069 | 0.855 |
| Hb-;Run+ | 0.06 | 0.065 | 0.845 |
| Hb-;Run- | 0.042 | 0.048 | 0.846 |
| Cad+;Kr- | 0.033 | 0.036 | 0.876 |
| Ftz+;Run- | 0.033 | 0.035 | 0.854 |
| Hb-;Kr- | 0.024 | 0.025 | 0.861 |
| Ftz+;Kr- | 0.017 | 0.018 | 0.889 |
| h+;Run- | 0.015 | 0.016 | 0.881 |
| Prd+;Run- | 0.01 | 0.01 | 0.868 |
| h+;Kr- | 0.007 | 0.009 | 0.882 |
| Kr-;Prd+ | 0.007 | 0.007 | 0.869 |
| Kr-;Tll- | 0.004 | 0.005 | 0.827 |

**Table 6:** PP17 interactions, filtered at level *τ* = 0.5 and ranked according to Δ*P*(*S*∣1).

| Interaction | $\Delta P(S\vert 1)$ | $P(S\vert 1)$ | $P(1\vert S)$ |
| --- | --- | --- | --- |
| Cad+;Tll+ | 0.376 | 0.384 | 0.859 |
| Cad+;Ftz+ | 0.153 | 0.157 | 0.876 |
| Cad+;Run+ | 0.141 | 0.145 | 0.875 |
| Cad+;Kni- | 0.099 | 0.101 | 0.846 |
| Cad+;D+ | 0.095 | 0.103 | 0.861 |
| Ftz+;Tll+ | 0.092 | 0.094 | 0.862 |
| Run+;Tll+ | 0.082 | 0.084 | 0.882 |
| Cad+;Ftz+;Tll+ | 0.08 | 0.08 | 0.897 |
| Ftz+;Hb+ | 0.076 | 0.078 | 0.893 |
| Hb+;Kr- | 0.074 | 0.076 | 0.893 |
| Cad+;Run+;Tll+ | 0.074 | 0.074 | 0.911 |
| Cad+;Kr-;Tll+ | 0.073 | 0.073 | 0.87 |
| D+;Tll+ | 0.069 | 0.07 | 0.879 |
| Cad+;D+;Tll+ | 0.06 | 0.06 | 0.893 |
| D+;Hb+ | 0.048 | 0.049 | 0.883 |
| Cad+;h+ | 0.047 | 0.049 | 0.848 |
| Cad+;Gt- | 0.046 | 0.048 | 0.824 |
| Cad+;Hkb- | 0.037 | 0.042 | 0.852 |
| h+;Tll+ | 0.031 | 0.031 | 0.846 |
| Hkb-;Tll+ | 0.026 | 0.027 | 0.88 |
| h+;Hb+ | 0.023 | 0.024 | 0.873 |
| D-;Kr- | 0.015 | 0.016 | 0.816 |
| D-;Ftz+ | 0.015 | 0.016 | 0.831 |
| D-;Run+ | 0.014 | 0.015 | 0.841 |
| h-;Kr- | 0.012 | 0.013 | 0.825 |
| Gt-;Kr- | 0.01 | 0.011 | 0.83 |
| Kr+;Run+ | 0.009 | 0.01 | 0.836 |
| Kni-;Tll- | 0.007 | 0.009 | 0.792 |
| Run+;Shn- | 0.006 | 0.007 | 0.834 |
| Hkb-;Kni- | 0.006 | 0.007 | 0.869 |
| Gt-;h+ | 0.003 | 0.004 | 0.806 |
| Hkb+;Tll- | 0.003 | 0.006 | 0.768 |
| Hb-;Kr+ | 0.002 | 0.011 | 0.727 |

**Table 7:**
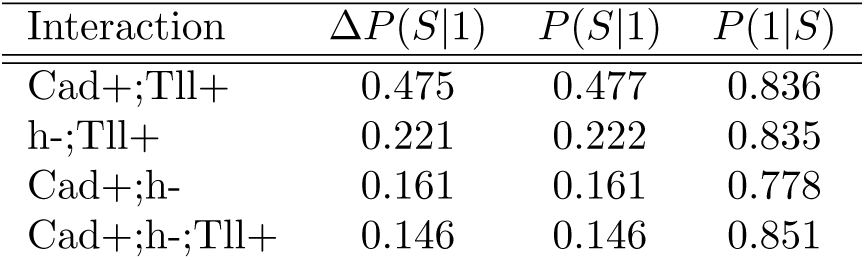
PP20 interactions, filtered at level *τ* = 0.5 and ranked according to Δ*P*(*S*∣1).

#### 5.2.1 Reconstructing the gap gene network through PP prediction

To determine whether signed interactions reflect known physical interactions, we evaluated our results in the context of the well-studied gap gene network. We constructed spatially local signed interaction networks (SLSIN) similar to the spatially local correlation networks (SLCN) studied in [37]. Following [37], we defined a given gene as expressed in PP region *i* if its corresponding PP coefficient *α_i_* ≥ 0.1. For each gap gene, we asked which interactions were predictive and stable for the PP(s) the gene is expressed in. More specifically, we filtered interactions from each PP prediction problem using algorithm 1 with *τ* = 0.5. We considered the interactions that remained after filtering to be candidate regulators for gap genes expressed in the predicted PP region. For gap genes expressed in multiple PPs, we considered contiguous domains separately as in [37].

Table 8 reports the known gap gene interactions described in [37, 19] by contiguous spatial domains. Genes on the left side of the arrow represent those whose spatial domain can be predicted by bracketed signed interactions on the right side of the arrow. We note that in all cases, several signed interactions involving gap genes are predictive of a particular spatial domain. We report the signed interaction with the highest value of Δ*P*(*S*∣1). Our analysis recovers all of the gap gene interactions reported in [19]. In addition, table 8 identifies known higher order relationships. For instance, *Caudal (Cad*) is known to play an interacting role with other gap genes in establishing the anterior posterior axis [19]. Signed interactions provide this additional insight that is missed when looking only at pairwise relationships.

**Table 8:** Gap gene interactions recovered from PP prediction using s-iRF. Each row corresponds to one link of the gap gene network in Fig. 7. For each pair of gap genes, we report the signed interaction containing Gene 2 (resp Gene 1) with the highest Δ*P*(*S*∣1) recovered when predicting the PPs of Gene 1 (resp Gene 2). This procedure recovers all links in the gap gene network with the correct sign.

| Link | Gene 1 | Gene 1 PP | Gene 2 | Gene 2 PP | Signed interactions |
| --- | --- | --- | --- | --- | --- |
| 1 | *Gt<sub>1</sub>* | 6, 7 | *Kr* | 8 | $Gt_1 \leftarrow \{Kni-, Kr-\}, Kr \leftarrow \{Gt-, Prd+\}$ |
| 2 | *Hb<sub>1</sub>* | 6-8 | *Kr* | 8 | $Hb_1 \leftarrow \{Kni-, Kr-\}, Kr \leftarrow \{Gt-, Hb-\}$ |
| 3 | *Hb<sub>1</sub>* | 6-8 | *Kni* | 8, 9 | $Hb_1 \leftarrow \{Kni-, Prd+\}, Kni \leftarrow \{Gt-, Hb-\}$ |
| 4 | *Kni* | 8, 9 | *Kr* | 8 | $Kni \leftarrow \{Cad+, Kr-\}$ |
| 5 | *Gt<sub>2</sub>* | 9 | *Kr* | 8 | $Gt_2 \leftarrow \{Cad+, Kr-\}, Kr \leftarrow \{Gt-, Prd+\}$ |
| 6 | *Gt<sub>2</sub>* | 9 | *Kni* | 8, 9 | $Kni \leftarrow \{Gt-, Prd+\}$ |
| 7 | *Hb<sub>2</sub>* | 17, 20 | *Kni* | 8, 9 | $Hb_2 \leftarrow \{Kni-, Tll-\}, Kni \leftarrow \{Gt-, Hb-\}$ |
| 8 | *Gt<sub>2</sub>* | 9 | *Hb* | 17, 20 | $Gt_2 \leftarrow \{D+, Hb-\}, Hb \leftarrow \{Cad+, Gt-\}$ |
| 9 | *Kni* | 8, 9 | *Tll* | 17, 20 | $Kni \leftarrow \{Cad+, Tll-\}, Tll \leftarrow \{Cad+, Kni-\}$ |
| 10 | *Gt<sub>2</sub>* | 9 | *Tll* | 17, 20 | $Gt_2 \leftarrow \{Cad+, Tll-\}, Tll \leftarrow \{Cad+, Gt-\}$ |
| 11 | *Hb<sub>2</sub>* | 17, 20 | *Hkb* | 20 | $Hb_2 \leftarrow \{Cad+, Hkb-\}$ |

## 6 Conclusion

Here we proposed signed iRF, or s-iRF, for refining the interactions recovered by iRF to gain insights into complex data. s-iRF builds on the notion of signed interactions, which describe both a set of interacting features and a functional relationship between features and responses. We validated our approach across multiple simulations and real data settings. In our simulations, s-iRF correctly identified active features as well as the functional relationship between features and responses in different regions of the feature space. For the problem of predicting enhancer activity, s-iRF recovered one of the few experimentally validated order-3 TF interactions and suggested novel enhancer elements where it may be active. Moreover, top four order-3 interactions we recover are among TFs with experimentall validated pairwise interactions, providing new candidate high-order interactions. In the case of spatial gene expression patterns, s-iRF recovers all 11 links in the gap gene network and suggests a high level of redundancy in *Drosophila* A-P patterning.

We have offered evidence that s-iRF extracts interpretable and relevant information from a fitted RF. However, there are several areas for further work. s-iRF searches for interactions globally despite the fact that iRF learns localized interactions in heterogeneous data. We are currently investigating strategies to search for interactions across meaningful subpopulations to gain additional insights into how iRF achieves accurate prediction. In addition, the signed interactions we recover represent individual components of the full iRF. We are exploring how these components relate to one another and how their combined behavior relates to responses to provide a more holistic representations of relationships in complex data.

Interpreting machine learning algorithms that achieve state-of-the-art predictive accuracy has the potential to offer new insights into high-dimensional, heterogeneous data. As part of this broad framework, the work we described here will help guide inquiry into the mechanisms underlying biological and other complex systems.

## Acknowledgments

This research was supported in part by grants NHGRI U01HG007031, ARO W911NF1710005, ONR N00014-16-1-2664, DOE DE-AC02-05CH11231, NHGRI R00 HG006698, DOE (SBIR/STTR) Award DE-SC0017069, DOE DE-AC02-05CH11231, and NSF DMS-1613002. SEC was supported by NIH grant NIH R01-GM076655. We thank the Center for Science of Information (CSoI), a US NSF Science and Technology Center, under grant agreement CCF-0939370. We thank T. Arbel for preparing *Drosophila* dataset.

## 7 Appendix

### 7.1 Interpretable prediction through signed interactions

RFs leverage high-order feature interactions for accurate prediction, but it is often difficult to interpret how these interactions influence predictions across an ensemble of trees. We use signed interactions to generate rule-based predictions that depend only on active features and that define a consistent functional relationship between features and responses. Specifically, for a signed interaction *S* we collect all leaf nodes throughout an RF for which *S* falls on the decision path. We represent this collection as a group of regions and predictions

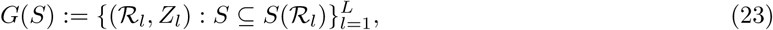

where 𝓡_*l*_ ⊆ ℝ^*p*^ and *Z_l_* ∈ {0, 1} denote the hyprerectangle and prediction associated with each leaf node *l* = 1 …, *L* in an RF. Equation (23) defines an ensemble of grouped rules, for which we write predictions as

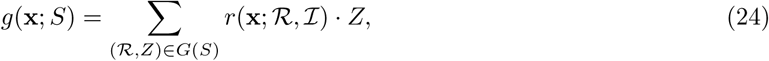

where 𝓘 = {∣ *j* ∣: *j* ∈ *S*} denotes the set of active features *S* is defined over and

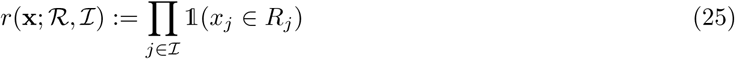

represents a rule over only the interacting features 𝓘. Predictions may assign equal weight to each term in equation (24) or weight based on measures of importance associated with each leaf node.

Intuitively, equation (24) can be viewed as a decision rule with smooth boundaries that exhibit several desirable properties.

1. Smooth decision boundaries make predictions less sensitive to small changes in **x** compared with a single decision rule defined over the same features.
2. Predictions incorporate non-linear feature interactions in the form of rules as in equation (25).
3. Predictions depend solely on the set of interacting features 𝓘.
4. Predictions are built from a collection of rules that define an identical functional relationship with responses described by (6).
5. Predictions are monotonic as a function of each active feature, and the sign of each *j* ∈ *S* defines whether predictions are increasing or decreasing as a function of the corresponding feature.

These properties ensure that *g*(**x**; *S*) provide robust and explainable predictions based on high-order, nonlinear interactions.

### 7.2 Region boundaries and response surfaces

Since signed interactions provide a coarse representation of hyperrectangles associated with a decision rule, running RIT on the collection of signed feature-index set/response pairs can be thought of as an approximate intersection of hyperrectangles. Understanding how response behavior varies across these coarse regions may be satisfactory in many applications. For instance, if the precision of experimental interventions is limited, a practitioner may not need to know the thresholds associated with each rule in a group. However, rule groups *G*(*S*) describe the distribution of thresholds selected across an RF, which we leverage to characterize response behavior with greater precision. This may be of interest in settings where a practitioner has the ability to precisely control some system and would like to optimize response behavior.

Intuitively, our representation uses each region 𝓡 ∈ *G*(*S*) as a data-adaptive binning to generate a histogram of response values. Averaging over every region 𝓡 ∈ *G*(*S*) allows us to identify the smoothed boundaries learned by an RF. We note that it may be desirable to weight regions 𝓡, for instance by using response purity or node size. In our simulations, we find that the boundary estimates are improved for larger leaf nodes and we weight each region in our response surface figures by the number of observations contained in the corresponding leaf. It may also be of interest to regularize estimates of these response surfaces, though we do not explore the idea in this paper.

When ∣*S*∣ is small we can visualize the response surfaces described in equation (24) (Fig. 5). For larger interactions, we visualize region boundaries marginally for each feature *j* ∈ 𝓘 by considering the distribution of thresholds that describe each *R_j_*, the 1-dimensional hyperrectangle corresponding to the *j^th^* feature in region 𝓡. From this perspective, variability in thresholds provides sense of the stability of boundaries under different data and model perturbations.

### 7.3 Redundant feature selection on decision paths

In some cases, particularly for deep decision trees used by RF, a feature *j* may be selected multiple times on the same decision path. In these situations both signed feature indices *j* and −*j* could appear on the decision path. The recursive nature of decision tree partitions suggests a natural way to handle this issue. For any features that are selected multiple times on a decision path, we use only the signed feature index associated with the first split, since subsequent splits on the same feature are restricted by the initial split. This representation destroys the one-to-one correspondence between a signed interaction and feature splits along a decision path. However, we find empirically that our simplified representation performs comparably to the full decision rule in terms of predictive accuracy and allows for a high-level, interpretable grouping of decision rules.

